# Hsp78-interaction proteome highlights chaperone role during protein aggregation in mitochondria under heat stress

**DOI:** 10.1101/2022.01.28.478199

**Authors:** Witold Jaworek, Marc Sylvester, Giovanna Cenini, Wolfgang Voos

## Abstract

Proteins of the Hsp100 chaperone family support protein homeostasis, the maintenance of protein activity under stress, by refolding aggregated proteins or targeting them for degradation. Hsp78, the ClpB-type mitochondrial member of the Hsp100 family, can be found in lower eukaryotes like yeast. Although Hsp78 has been shown to contribute to protection against elevated temperatures in yeast, the biochemical mechanisms underlying this mitochondria-specific thermotolerance are still largely unclear. To identify endogenous chaperone substrate proteins, we generated an Hsp78-ATPase mutant with a stabilized substrate binding behaviour. We used two SILAC-based quantitative mass spectrometry approaches to analyze the role of Hsp78 during heat stress-induced mitochondrial protein aggregation and disaggregation processes on a proteomic level. In the first setup, Hsp78-interacting polypeptides were identified to reveal the endogenous substrate spectrum of the chaperone. Our analysis revealed that Hsp78 is interacting with a wide variety of proteins related to metabolic functions including energy production and protein synthesis, as well as other chaperones, thus maintaining crucial functions for mitochondrial stress resistance. We compared these interaction data with a second experimental setup that focussed on the on overall aggregation and disaggregation processes in mitochondria under heat stress on a proteomic level. This revealed specific aggregation-prone protein populations and demonstrated the direct quantitative impact of Hsp78 on stress-dependent protein solubility under different conditions. We conclude that Hsp78 together with its cofactors represents a recovery system that protects major mitochondrial metabolic functions during heat stress as well restores protein biogenesis capacity after return to normal conditions.

## Introduction

Every organism requires specific biochemical processes for the maintenance of the internal protein homeostasis, keeping cellular functions intact and the cell alive. Due to the exposition to variable environmental conditions, especially immobile organisms like bacteria, fungi and plants suffer from thermal stress and therefore exhibit distinct protective mechanisms that convey a certain degree of thermotolerance. Also human cells may be subjected to heat stress under certain pathological conditions like fever. These proteo-protective mechanisms are based on the activity of molecular chaperones, representing ATP-dependent enzymes that stabilize misfolded proteins, protecting them from aggregation and support degradation processes (1-4). One important group of chaperones with a major role during protein aggregation reactions is the Hsp100 chaperone family. They belong to the AAA+ class of enzymes (ATPase associated with wide variety of cellular activities), many of them involved in molecular remodelling reactions (5). Enzymes of a particular subgroup of the Hsp100 family, all relatives of the bacterial ClpB protein, are able to resolve polypeptides from insoluble aggregates and to support their refolding via the Hsp70 chaperone system or their degradation through various ATP-dependent proteases (6,7). ClpB-type chaperones typically form hexameric ring structures that are formed by intermolecular interactions of their two AAA protein domains. It was proposed that the mechanistic basis of the chaperone activity is based on a threading mechanism of the polypeptide chains through the central pore of the ring complex, thereby dissolving any incorrect protein conformations (3). Bacterial ClpB was shown to cause thermal hypersensitivity in the respective null-mutant strains, thus ClpB-type chaperones facilitate thermal tolerance to bacteria (8). In comparison to other Hsp100 chaperones, ClpB-type proteins do not directly assemble with proteases (9).

The problematic nature of heat stress might be even more relevant for mitochondrial proteins as it is assumed that, due to metabolic processes, temperature inside the organelle may be elevated permanently (10). The basic biochemical mechanisms of the chaperone enzymes involved in mitochondrial aggregation have been largely established to date, either based on the utilization of artificial reporter proteins or the characterization of the thermal reactivity of single examples of specific proteins. However, for a complete understanding of mitochondrial protein homeostasis under thermal stress conditions, it is fundamental to identify proteins that exhibit less stable folding states and have a tendency to aggregate. Previous experiments exposing human cells to mild heat stress, similar to fever conditions, exhibited an overall quite robust behaviour of mitochondrial proteins with only few polypeptides prone to aggregation (11). In contrast, a study in yeast established a strong effect of elevated temperatures on mitochondrial morphology (12). Yeast cells contain two homologs of bacterial Hsp100 family chaperone ClpB, the cytosolic Hsp104 (13), and the mitochondrial Hsp78 (14). As ClpB family members are not present in higher eukaryotes (15), it has been proposed that other chaperones, in particular mtHsp70 together with its co-chaperone Tid1, may take over Hsp100 function in human mitochondria (16). Similar to other Hsp100 family members, Hsp78 is an ATP-dependant enzyme, which is up-regulated upon thermal stress and most likely forms the typical barrel-shaped hexameric enzyme complexes with a central pore (17,18). In analogy to bacterial ClpB, it is assumed also for Hsp78 that substrate polypeptides are mechanically pulled through the pore and are thereby disaggregated for further processes executed by other chaperone systems (19). Similar to other members of the family, Hsp78 contains two AAA-type nucleotide binding-domains with conserved Walker A and Walker B motifs (20), which are connected by a coiled-coil middle segment (M domain) that is proposed to provide an interaction with Hsp70-type chaperones, stimulating ATPase function (21). Due to its organellar localisation, Hsp78 contains an N-terminal mitochondrial targeting sequence but lacks the N-terminal substrate-binding domain of bacterial ClpB (17). It is assumed that the N-terminal domain of ClpB plays a role in disaggregation of large aggregates (22), while there is little known about the mechanism of substrate interaction in case of the mitochondrial version.

Knowledge about the proteome stability inside mitochondria and the substrate patterns of the Hsp78 chaperone would provide fundamental information for the understanding of pathological patterns related to mitochondrial stress. So far, although some information is available on a whole-cell level (23), no comprehensive proteomic study was done in yeast mitochondria with the focus on Hsp78 disaggregation activity after heat stress. In our new study, we constructed a mutant form of Hsp78 with both AAA domains mutated in their Walker B motifs (Hsp78_TR_). We aimed on generating a stabilized substrate-chaperone interaction due to an impaired capability to hydrolyse ATP. A similar approach was pursued for bacterial ClpB to identify endogenous substrate proteins (24). We performed a basic biochemical characterisation of the mutant enzyme in order to check characteristics like complex formation and ATPase activity, as well as its general protein disaggregation competence. We focused on the identification of Hsp78-interacting proteins under normal and stress conditions, utilizing a quantitative proteomics approach. In a second quantitative mass spectroscopy (qMS) experiment, we demonstrate differences in protein aggregation and aggregate recovery between WT and mutant Hsp78 inside mitochondria on a proteomic level. The analysis revealed that Hsp78 plays a critical role in mitochondrial protein homeostasis with a focus on the reactivation of essential mitochondrial functions like TCA cycle and the respiratory chain but also including protein synthesis and import, after thermal stress exposure.

## Material and Methods

### Plasmids and strains

Constructs used in this study were generated by conventional cloning procedures. Hsp78_TR_ mutant was generated from plasmid pTB25by side directed mutagenesis (SDM) in two steps. First E216Q was inserted by SDM creating plasmid pWJ01. In the second step pWJ02 was created by SDM inserting E612Q. Subsequently the construct was cut out by BamHI and AvrII and ligated into pTB10 resulting in a construct without histidin-tag (pWJ04). In the last step, plasmid sequence was determined and a point mutation (D386M) was recovered by SDM creating pWJ05. For tagged constructs the mutated segment of Hsp78_TR_ was cut with BamHI and AvrII and inserted into pKR09 (Hsp78-6xHis) replacing the wild-type segment creating pWJ07. In both pYES plasmid constructs the encoded Hsp78 variants were under control of the galactose-inducible promotor GAL10/1. As control, expression via the plasmid pHSP78 was used to represent the untagged version.

All experiments were performed with *S. cerevisae* strain Y03617-ES except in the qMS experiments, which were exclusively performed with Y13617-ES due to the use of lysine isotope forms. A list of all plasmids and strains used in this study is attached in table S1.

### Expression of Hsp78 constructs in yeast cells

Yeast cells were inoculated over night in suitable selective growth medium containing 3% raffinose and grown at 30°C. Cells were transferred to a medium with 3% ethanol as carbon source for respiratory growth. Expression was induced at OD 1 by addition of 2% galactose. Constructs were expressed for up to 4 hours. For total cell extracts cells were reisolated and solved in 50 μl H_2_0. 250 mM NaOH and 1,2% beta-mercaptoethanol was added. After an incubation for 10 min on ice cells were TCA precipitated and 0.2 OD was loaded per lane for 12.5% SDS-PAGE.

### In vitro ATPase assay

Hsp78 ATPase activity was measured by a coupled reaction for enzymatic phosphate release by PiColorLock Gold Phosphate Detection System (Novus Biologicals, Wiesbaden, Germany). For comparison a standard curve of phosphate (0-50 μM) was measured. WT Hsp78 and mutant Hsp78_TR_ expressed and affinity purified from *E.coli* were diluted to a final concentration of 0.08 μg/μl in a volume of 500 μl reaction buffer (80 mM KCl, 10 mM MgCl_2_, 1 mM DTT, 0.25 mM ATP, 30 mM Tris, pH 7.4). Experiments were incubated at 37°C and 90 μl samples were collected every 5 min up to 20 min of reaction time. Control samples were only determined after 20 min. Subsequently enzymatic reaction Gold mix was added (1:5) to the mixture. After a short incubation of 5 min at RT, 10% of stabilizer was added and the samples were incubated for 30 min at RT according to manufactures directions. The resulting phosphate-complex was quantified by the absorbance at 635 nm in a TECAN plate reader (96-well format).

### Growth and acquired thermo tolerance assays

Yeast cells were inoculated over night in suitable selective growth medium containing 3% raffinose and grown at 30°C. Cells were transferred to a medium with 3% ethanol as carbon source for respiratory growth. Expression was induced at OD 1 by sedimentation and resuspension in 2% galactose containing medium. Cells were induced for 3 h and then heat-treated at 42°C for 60 min in a water bath with a subsequent regeneration step for 30 min at 30°C and a final heat exposition at 48°C for 60 min. 10^6^ cells were then diluted with water in 1:3 dilution steps and 7 μl cell suspension was spotted on selective growth plates. Plates were containing either 2% glucose for control or 3% glycerol for analysis of the mitochondrial fitness and incubated at 30°C for several days.

### Hsp78 complex formation by Blue Native-PAGE

Intra-mitochondrial Hsp78 complex formation was analyzed by BN-PAGE. For that 100 μg isolated mitochondria were incubated in buffer (250 mM sucrose, 80 mM KCl, 10 mM MOPS/KOH, pH 7.2) containing an ATP regeneration system (10 mM KPi, 10 mM creatine phosphate, 5 mM MgCl_2_, 4 mM NADH, 3 mM ATP, 50 μg/ml creatine kinase) or buffer depleted for their nucleotide content (1 mM EDTA, 0,01 U/μl apyrase) for 10 min at RT. After reisolation mitochondria were lysed in 100 μl ice-cold digitonin lysis buffer (50 mM KCl, 20 mM Tris/HCl, pH 7.4, mM EDTA, 1 mM PMSF, 10% glycerol, 1% digitonin) by pipetting (25x). For gradient BN-PAGE (5-15%) 70 μl per lane was loaded while for SDS-PAGE control (12,5%) 30 μl was loaded. In addition protein complex formation was also analyzed for purified Hsp78 proteins expressed in *E. coli.* To assay complex stability, 4 μg of protein was diluted in a final volume of 60 μl in protein buffer L (50 mM KCl, 30 mM HEPES, pH 7.4, 5 mM MgCl_2_, 4 mM ATP, 2 mM EDTA, 20% glycerol) or protein buffer H (400 mM KCl, 30 mM HEPES, pH 7.4, 2 mM EDTA, 20% glycerol). After addition of 6 μl 10x BN loading dye whole samples were used. BN-PAGE was performed according to a procedure elaborated for the analysis of multi-protein complexes from cellular lysates (25).

### Aggregation and recovery assays

The [^35^S]-radiolabeled preprotein cytb_2_(107)-DHFR_DS,_ generated using the TNT T7 Quick Coupled Transcription/Translation System (Promega), was imported into isolated mitochondria for 20 min at RT and import was stopped by the addition of 0.5 μM valinomycin. Successful and complete import was controlled by resistance of the imported polypeptides against treatment with externally added proteinase K (26). Assessment of aggregation and recovery of the imported protein were performed according to an established procedure (27) with 5 min heat stress and 60 min recovery. For the analysis of the aggregation tendency of mitochondrial proteins, isolated mitochondria were either maintained at RT, stressed for 20 min at 42°C or stressed and recovered for 60 to 120 min. Three times 200 μg mitochondria from *hsp78Δ* cells were treated at 45°C for 20 min to produce aggregates without attached Hsp78. Mitochondria were lysed by pipetting (20x) in 800 μl lysis buffer (200 mM KCl, 30 mM Tris/HCl. pH 7.4, 4 mM NADH, 3 mM ATP, 0.5 mM PMSF, 0.5% Trition X-100, protease inhibitor) each. Samples were centrifuged for 30 min at 4°C and 125,000 xg to produce aggregate pellets. In parallel, mitochondrial lysates were obtained from 120 μg isolated mitochondria (see above) that expressed Hsp78_WT_ or Hsp78_TR_ proteins. After centrifugation for 30 min at 4°C and 125,000 xg 160 μl of each cleared Hsp78 lysate corresponding to 40 μg of mitochondria was put on the *hsp78Δ* aggregate pellets corresponding to an amount of 200 μg of mitochondria. Samples were incubated at 30°C for 30 min to allow substrate binding of Hsp78. Finally, the samples were centrifuged again and supernatant and pellet fractions were collected. While all supernatants taken were directly TCA precipitated, pellet fractions were first washed with 150 μl SEM buffer (250 mM sucrose, 10 mM MOPS, 1 mM EDTA, pH 7.2). Mock samples were added for all reactions. Protein bands were detected after SDS-PAGE by digital autoradiography and quantified by Multi Gauge software (Fujifilm) or by Western blot using specific antibodies.

### Size exclusion chromatography

Size exclusion chromatography was performed on Äkta purifier system (Cytiva, Freiburg, Germany) with HiLoad 16/600 Superdex 75 column. For the run, 1 mg of Ni-NTA affinity purified protein expressed in *E. coli* was used. Buffer GFL (50 mM KCl, 30 mM HEPES, pH 7.4, 5 mM MgCl_2_, 2 mM ATP, 2 mM DTT, 2 mM EDTA, 20% glycerol) or buffer GFH (400 mM KCl, 30 mM HEPES, pH 7.4, 2 mM DTT, 2 mM EDTA, 20% glycerol) were used to dilute protein solution to an injection volume of 500 μl. Same buffers were used for the equilibration of the column and the run. Molecular weight markers were run only with buffer GFL. For Western blot analysis of the samples 0.5 ml fractions were collected.

### Analysis of Hsp78-interacting proteins via qMS

Yeast cells cultured in minimal medium were incubated with different isotope forms of lysine (SILAC, stable isotope labelling with amino acids in cell culture). For cells expressing non-tagged Hsp78_WT_ normal lysine was used, for Hsp78_WT_-6xHis ^13^C_6_-^15^N_2_ L-lysine (heavy; +8) and for the mutant Hsp78_TR_-6xHis ^2^H_4_ L-lysine (medium; +4) was added, respectively. After purification from the cells, mitochondria were treated with different temperature conditions (see above). Mitochondria (1 mg) were lysed with 2 ml lysis buffer SA (80 mM KCl, 50 mM NaPi, pH 7.8, 5 mM MgCl2, 2 mM ATP, 0.5 mM PMSF, 5% glycerol, 0.5% Triton X-100, 1x protease inhibitor) and shaking for 10 min at 4°C. After centrifugation for 5 min at 12,000 x g and 4°C, supernatants were added to 50 mg Ni-TED column material that was previously washed with lysis buffer SA. Slurry was incubated rotating at 4°C for 15 min, and applied on mini-columns. The columns were washed three times with 1 ml lysis buffer SA each and three times with 1 ml buffer SW (80 mM KCl, 50 mM NaP_i_, pH 7.8, 5 mM MgCl_2_, 2 mM ATP, 0.5 mM PMSF, 5% glycerol, 1x protease inhibitor). Finally, proteins were eluted in three fractions with 1 ml elution buffer SE (250 mM imidazole, 80 mM KCl, 50 mM NaP_i_, pH 7.8, 5 mM MgCl_2_, 2 mM ATP, 0.5 mM PMSF, 5% glycerol, 1x protease inhibitor) each. Subsequently, similar eluates volumes containing the three different Hsp78 forms were pooled and TCA-precipitated and run together on a SDS-PAGE gel. For MS sample preparation, gel pieces were cut after Coomassie G-250 staining. The amount of purified bait protein served as internal loading control while the untagged Hsp78 eluates defined the non-specific background signals.

### Analysis of mitochondrial protein aggregation and recovery qMS

For the *in organello* aggregation and recovery assay isolated mitochondria were loaded on a multi step sucrose gradient to ensure high sample purity with low contaminations from other cellular components. The gradient was built by three layers with 4 ml 32%, 1.5 ml 23% and 1.5 ml 15% sucrose (w/v) solved in EM buffer (10 mM MOPS/KOH, pH 7.2, 1 mM EDTA). Mitochondria were added on top of the gradient and covered with 1.5 ml EM buffer. After centrifugation at 125,000 x g and 4°C for 60 min in a Beckmann SW41 Ti swinging-bucket the mitochondria were re-extracted from the tube bottom in re-suspension buffer (250 mM sucrose, 80 mM KCl, 10 mM MOPS/KOH, pH 7.2, 5 mM MgCl_2_). Mitochondrial content was measured by Bradford assay and adjusted to a concentration of 200 μg/ml in resuspension buffer. Differently labelled mitochondria (see co-purification SILAC experiment) were mixed in a 1:1 ratio using 50 μg each. Mitochondria expressing untagged Hsp78 were excluded. Samples were treated similarly to Aggregation and recovery experiments. For MS sample preparation, gel pieces were cut after SDS-PAGE.

### Peptide preparation

Gel slices were subjected to tryptic in gel digestion (28). In brief, slices were washed consecutively with water, 50% and 100% acetonitrile (ACN). Proteins were reduced with 20 mM DTT in 50 mM ammonium bicarbonate and alkylated with 40 mM acrylamide (in 50 mM bicarbonate). The slices were washed again and dehydrated with ACN. Dried slices were incubated with 250 ng sequencing grade trypsin at 37°C overnight. The peptide extract was separated and remaining peptides extracted with 50% ACN. Peptides were dried in a vacuum concentrator and stored at −20°C.

### LC-MS measurements of peptides

Peptides were dissolved in 10 μl 0.1% formic acid (FA) and 3 μl were injected onto a C18 trap column (20 mM length, 100 μm inner diameter, ReproSil-Pur 120 C18-AQ, 5 μm; Dr. Maisch GmbH, Ammerbuch-Entringen, Germany) made in-house. Bound peptides were eluted onto a C18 analytical column (200 mM length, 75 μm inner diameter, ReproSil-Pur 120 C18-AQ, 1.9 μm, with 0.1% formic acid as solvent A). Peptides were separated during a linear gradient from 2% to 35% solvent B (90% acetonitrile, 0.1% FA) within 83/90 min at 250 nl/min. The nanoHPLC was coupled online to an LTQ Orbitrap Velos mass spectrometer (Thermo Fisher Scientific, Bremen, Germany). Peptide ions between 330 and 1600 m/z were scanned in the Orbitrap detector with a resolution of 60,000 (maximum fill time 400 ms, AGC target 10^6^). The 22 most intense precursor ions (threshold intensity 3000, isolation width 1.2/1.0 Da) were subjected to collision induced dissociation (CID, normalized energy 35) and analyzed in the linear ion trap. Fragmented peptide ions were excluded from repeat analysis for 20/15 s.

### MS data analysis

For peptide analysis raw data processing and database searches were performed with Proteome Discoverer software 2.1.1.21 (Thermo Fisher Scientific). Peptide identifications were done with an in-house Mascot server version 2.5.1 (Matrix Science Ltd, London, UK). MS2 data were searched against *Saccharomyces cerevisiae* sequences in SwissProt (release 2017_10). Precursor ion m/z tolerance was 8 ppm, fragment ion tolerance 0.5 Da. Tryptic peptides with up to two missed cleavages were searched. Propionamide on cysteines was set as a static modification. Oxidation of methionine, labels ^13^C_6_^15^N_2_, and ^2^H_4_ on lysine were allowed as dynamic modifications. Mascot results were assigned q-values by the Percolator algorithm (29) version 2.05 as implemented in Proteome Discoverer. Spectra with identifications below 1% q-value were sent to a second round of database search with semitryptic enzyme specificity (one missed cleavage allowed) and 10 ppm MS1 mass tolerance (propionamide dynamic on Cys). In co-purification experiments, only proteins were included if at least two peptides were identified with <1% FDR. Typical FDR values were ≤1% (peptide spectrum matches), 1.2% (peptides), and <1% (proteins). Only unique peptides were included in protein quantification. The mass spectrometry proteomics data have been deposited to the ProteomeXchange Consortium via the PRIDE (30) partner repository with the dataset identifiers PXD030405 (part A: protein interactions) and PXD030436 (part B: aggregation).

### Further data processing

In the co-purification experiments all abundance values obtained in the control mitochondria expression non-tagged Hsp78, representing non-specific background signals, were substracted from the respective values obtained with the tagged variants in the individual experiments and then an average value of 3 experiments was generated. In order to mathematically allow the calculation of co-purification enrichment ratios also in cases where the respective protein has not been identified in the background control sample, abundance values were set to a minimal abundance value of 10.000, representing background intensity. All protein abundances above background were corrected by Hsp78 bait protein abundance in the respective experiment. To assess the reliability of the qMS protein identification each protein hit was categorized concerning their identification reproducibility in isotope channels and replicate experiments. Category A proteins (best candidates) were identified with at least three peptides in all isotope channels and all experiments, category B in two replicates and category C in one of three replicates. Category D was not detected in any replicate of the respective temperature conditions but in others. Calculations were performed with Microsoft Excel and visualisation was done using GraphPad PRISM 6. Additional protein information was collected from the UniProtKB database (https://www.uniprot.org/). Functional categories were obtained using the STRING (Protein-Protein Interaction Networks) database (https://www.stringdb.org/).

### Miscellaneous

All chemicals were analytical grade and supplied from Carl Roth (Karlsruhe, Germany) if not indicated otherwise. Antisera used in this study were previously verified by band pattern comparisons of wild-type and respective knockout strains in *S. cerevisiae*. Enzymes were purchased from Thermo Fischer Scientific (Darmstadt, Germany). Chemicals for qMS analysis were from Sigma-Aldrich (Taufkirchen, Germany).

## Results

### Hsp78_TR_ shows altered ATPase turnover, but functional complex formation

We generated an Hsp78 ATPase-deficient mutant protein with both glutamic acids in the Walker B motifs exchanged to glutamine (E216Q/E614Q). Based on observation with a similar mutant form of the bacterial ClpB (24), we expected that the Hsp78 mutant would exhibit a higher substrate affinity to support the determination of the endogenous polypeptide substrate spectrum. The E216Q/E614Q mutant protein is abbreviated in the following as Hsp78_TR_. In order to facilitate protein purification, we added a C-terminal 6x polyhistidine-tag (Fig. 1A). To serve as control for the biochemical properties of the mutant protein, a similar wild-type (WT) version was also generated. Both constructs were inserted in a yeast expression plasmid that allowed expression the protein in a galactose-inducible fashion to avoid a long-term adaptation to the mutant phenotype. Proteins were expressed in a deletion background lacking the endogenous Hsp78 (*hsp78Δ*) so only the properties of the expressed proteins were analyzed. To check whether protein levels of wild-type and mutant proteins were comparable, we performed expression tests with both tagged proteins compared to an untagged WT version (Fig. 1B). Mutant Hsp78_TR_ was expressed slightly weaker than the corresponding WT version although the His-tag had no severe effect on the expression level (Fig. 1B). Furthermore, a thermotolerance growth assay on non-fermentable carbon source (glycerol) was performed to analyse mitochondrial functionality after heat pre-treatment at 42°C for 60 min and a subsequent thermal shock at an otherwise lethal temperature of 48°C for 60 min. Under these thermal stress conditions, the dependence of mitochondrial activity on the function of Hsp78 was clearly visible, showing a higher thermal resistance for cells expressing wild-type protein than mutant Hsp78_TR_. In contrast, growth on fermentable carbon source (glucose) was similar (Fig. 1C). ATPase activities of the Hsp78 variants were measured using purified protein expressed in *E.coli* by a coupled enzymatic assay. The assay revealed a strong ATPase impairment of 70% in the Hsp78_TR_ mutant compared to WT (Fig. 1C). In order to test complex formation of the expressed mutant Hsp78 we performed a blue-native (BN) PAGE with isolated mitochondria expressing the tagged constructs as described above. Higher molecular weight complexes containing Hsp78 were detected by Western blot with specific anti-Hsp78 antibodies. The majority of the Hsp78 signals were detected at sizes corresponding to a trimeric and hexameric complex with implications of even higher sizes for Hsp78_TR_ under ATP regenerating conditions (Fig. 1E). Moreover, we analyzed the purified proteins by BN-PAGE and size exclusion chromatography showing formation of similar complexes with additional smaller and even higher complexes. We also investigated complex stability by its resistance to high salt treatment. We found hsp78_TR_ being stable up to concentrations of 400 mM NaCl in the buffer, while wild-type protein was dissociating into smaller oligomers (dimers and trimers) (Fig. 1F). We concluded that a higher complex stability for the mutant likely reflected a stronger substrate binding capacity. Taken together, we could show that our Hsp78 mutant is basically functional but exhibited a reduced ATPase activity and a stabilized complex formation.

**Fig. 1:**
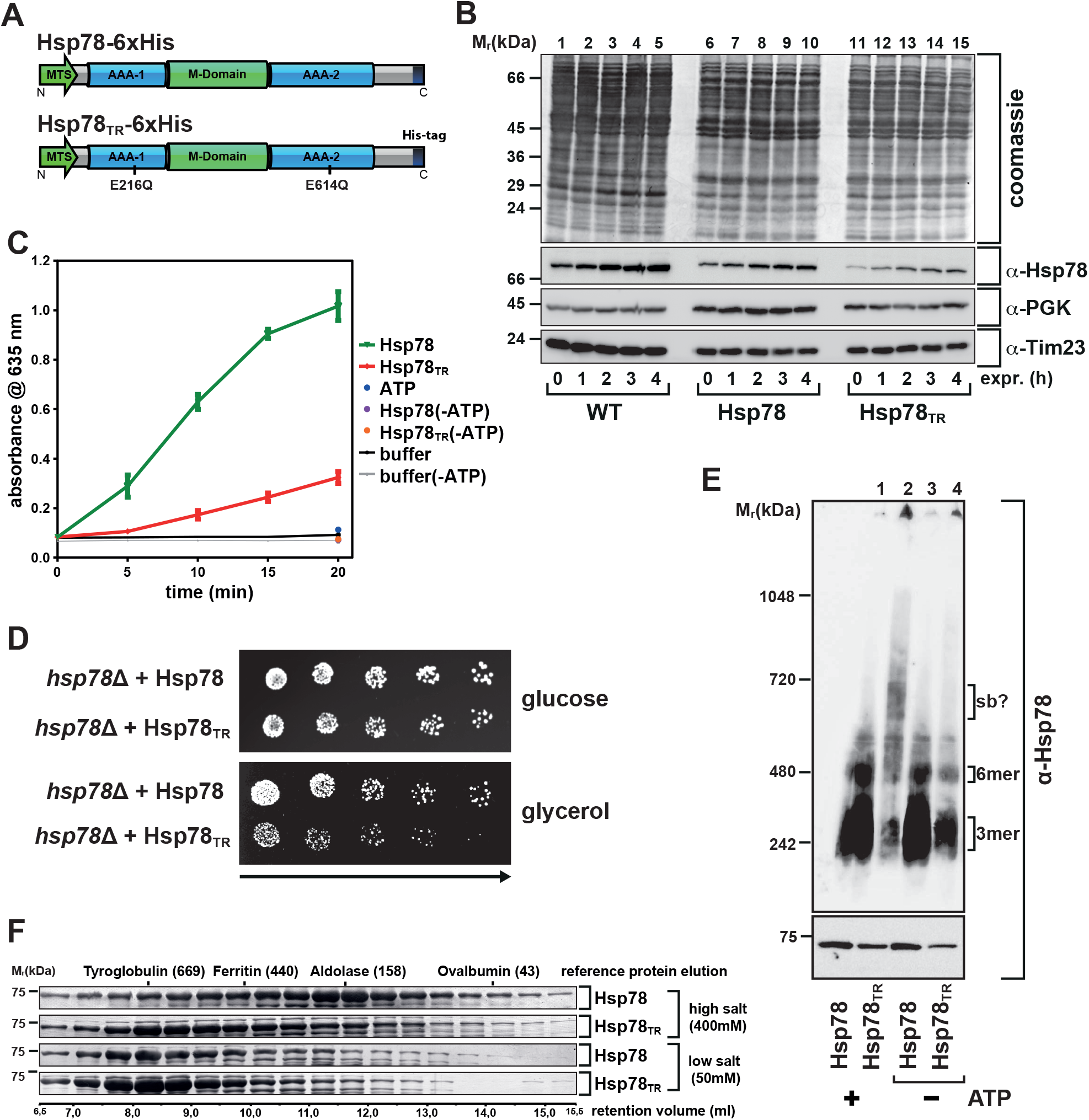
Hsp78 WalkerB mutations impair ATPase activity and lead to altered protein behavior. A, Schematic depiction of the WT (Hsp78) and mutant (Hsp78TR) protein tagged with six histidine residues used in this study. In the mutant both AAA-domains are modified in the Walker B section by amino acid exchange from glutamic acid to glutamine. B, Yeast hsp78Δ cells expressing non-tagged WT Hsp78 or the tagged proteins Hsp78/Hsp78TR were induced by addition of 2% galactose for the indicated time points and protein levels were detected by SDS-PAGE and Western blot using α-Hsp78 antiserum. Decoration with α-Tim23 for control of mitochondrial content and α-PGK for general loading was performed. The upper panel shows the blot membrane stained with Coomassie Blue. C, ATPase activity determination of Hsp78 (green curve), Hsp78TR (red curve) and control reactions as indicated. Proteins were expressed and purified from E. coli. Shown are mean values of three experiments; error bars indicate SEM. D, Serial dilution growth test for acquired thermotolerance of hsp78Δ cells expressing either Hsp78 or Hsp78TR with 1:3 dilution steps. Cells were either plated on glucose (control) or on glycerol containing plates and grown for at least 48 h at 30°C. E, BN-PAGE (upper panel) and control SDS-PAGE (lower panel) with corresponding Western blots decorated with α-Hsp78 using mitochondria isolated from of hsp78Δ cells expressing either Hsp78 or Hsp78TR. Mitochondria were resuspended either in resuspension buffer (+ATP) or in inhibition buffer (-ATP) prior to lysis. F, Sepharose based gelfiltration size exclusion chromatography with Hsp78 and Hsp78TR expressed and purified from E. coli. Fraction samples (0,5 ml each) were loaded on 12,5% SDS-PAGE and subsequent Coomassie Blue staining was performed. Protein samples were pre-incubated in high or low salt buffer as indicated. Retention volume indicates protein complex size by comparison with known high molecular proteins and complexes as reference.

### Hsp78 is the key mitochondrial disaggregase and binds actively to aggregated proteins

Furthermore, we tested the influence of the novel Hsp78 mutant proteins on the mitochondrial disaggregation reaction. As an aggregation reporter we utilized the fusion protein b_2_(107)Δ-DHFR_DS_ that contains a thermo-labile dihydrofolate reductase (DHFR) domain with a matrix-targeting mitochondrial presequence (27). This protein was imported in radioactively labelled form into isolated WT or mutant mitochondria. Aggregation of this protein was triggered by applying a short heat stress treatment of 42°C for 5 min. Afterwards, the resolubilization of this substrate during a recovery period of 20 min at 25°C was determined by a mild detergent lysis of the mitochondria, high-speed centrifugation of the lysate and the comparison of the radioactive signal intensities in pellet and soluble fractions. Experiments were conducted for isolated mitochondria expressing wild-type protein, overexpressed Hsp78, overexpressed Hsp78_TR_ and lacking Hsp78 (*hsp78Δ*). Results showed a strongly reduced recovery activity for Hsp78_TR_ with comparable values to knockout or wild-type control mitochondria without ATP supply (Fig. 2A-C). This shows that the inability of the mutant to hydrolyse ATP has a direct effect on its potential to resolubilize aggregated polypeptides.

**Fig. 2:**
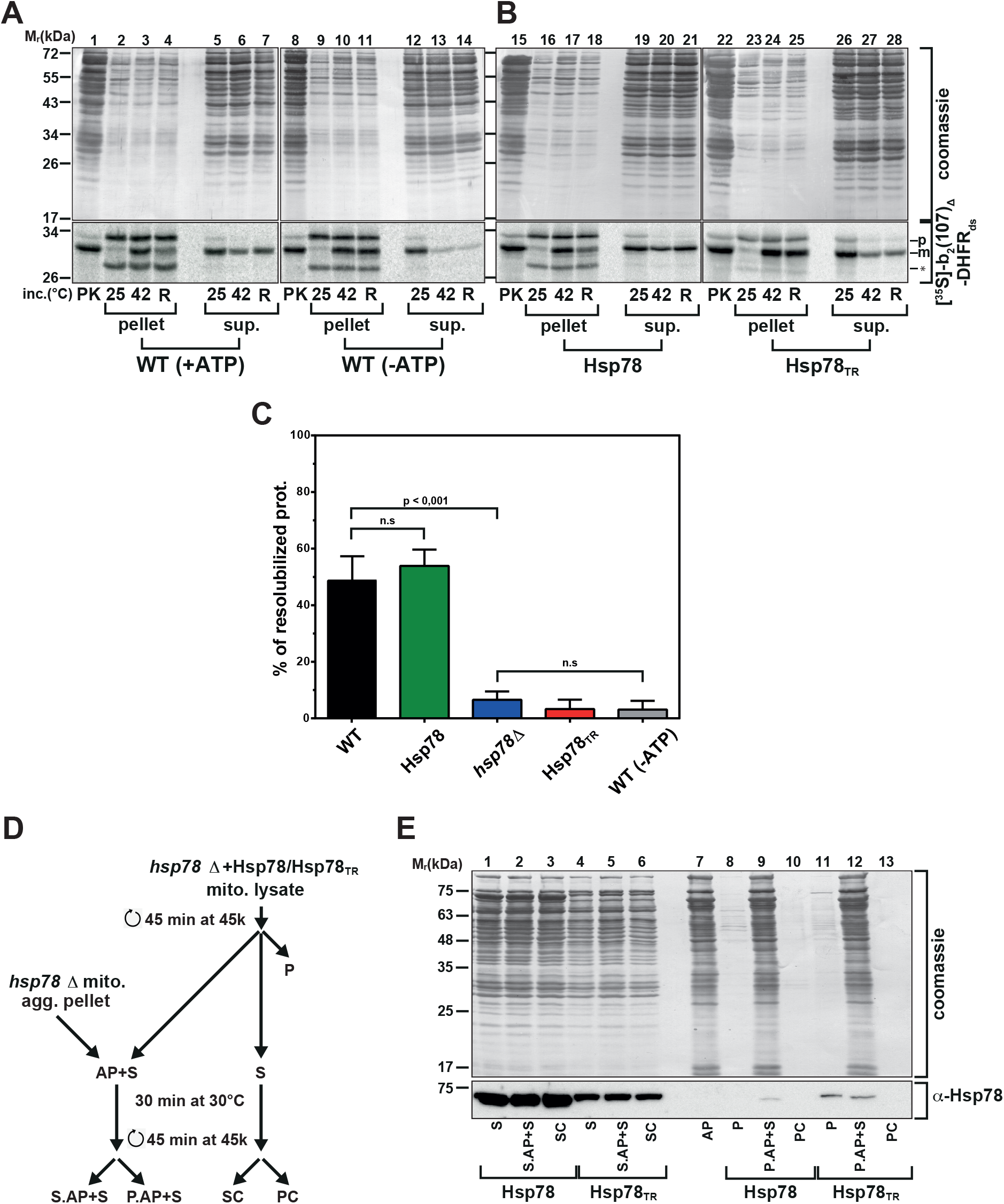
Hsp78 supports active disaggregation of heat induced mitochondrial protein aggregates. A, Disaggregation of imported reporter protein. [35S]-labeled b2(107)Δ-DHFRds was imported for 20 min at 25°C into isolated WT mitochondria. Mitochondria were then resuspended in presence (+ATP) or absence of ATP (-ATP) and kept at 25°C or stressed for 5 min at 42°C. After recovery at 25°C for 60 min (R), lysis and ultracentrifugation was performed for all samples. Supernatants and pellet fractions were collected and analysed by SDS-PAGE. Non-imported preproteins were digested by protease K as indicated (PK) preprotein (p) and mature form (m). Asterisk indicates additional translation product that has no influence on protein import. As loading control, the Coomassie Blue stained blot membrane is shown. B, Import was performed as described in A into mitochondria over-expressing tagged Hsp78 or Hsp78TR. Both samples were resuspended in presence of ATP and the same experimental procedure was applied as in A. C, Quantification of aggregation and recovery experiments as in A and B with additional inclusion of hsp78Δ mitochondria. Shown are bar diagrams with SEM and significance (n=5-7; n=2 (-ATP); n. s., non-significant). D, Flow scheme of aggregate binding assay (AP, aggregate pellet; S, high-speed supernatant of detergent-lysed mitochondria; C, control). Heat treatment and centrifugation conditions as indicated E, Interaction of Hsp78 variants with aggregated mitochondrial proteins. Incubation of an aggregate pellet derived from hsp78Δ mitochondria with the high speed centrifugation supernatants from mitochondria over-expressing tagged Hsp78 or Hsp78TR (as described in D) followed by SDS-PAGE, Western blot and decoration with α-Hsp78 (Coomassie Blue staining as loading control).

To elucidate substrate interaction behaviour of the Hsp78_TR_protein in comparison to the WT protein, we performed an aggregate-binding assay (experimental outline see Fig. 2C). First a mix of aggregated mitochondrial polypeptides, serving as potential Hsp78-substrates, was produced by a high-temperature treatment of *hsp78Δ* mitochondria and high-velocity centrifugation to separate soluble and insoluble protein populations, obtaining an aggregate pellet (AP) of mitochondrial proteins excluding any effect of Hsp78. Second, we generated cleared lysates of Hsp78 and Hsp78_TR_ expressing mitochondria (S) that were not subjected to any heat treatment as chaperone sources. Next the mitochondrial lysates containing the Hsp78 variants were incubated with the previously obtained resuspended aggregate pellet to monitor potential substrate binding reactions. Lysates and aggregated proteins were incubated together for 30 min at 30°C and then centrifuged again. The chaperone detected in the pellet of the second centrifugation therefore reflected the amount bound to aggregated polypeptides. As control for a potential intrinsic aggregation propensity of the mutant Hsp78_TR_, also the pellets (P) of mitochondrial lysates without any heat treatment were analyzed showing that only a minor amount of the mutated Hsp78_TR_ was pelleted, most likely because the mutations introduced a certain instability of the polypeptide structure. Compared to the total amount of chaperone added to the binding reaction, only a small amount was recovered in the aggregate pellet under these conditions. Although the expression of WT Hsp78 in the cleared lysates was higher than of Hsp78_TR_, the experiment showed that Hsp78_TR_ did co-sediment stronger with the heat-induced aggregates than Hsp78 (P.AP+S lanes, Fig. 2E). The equally treated control samples in the absence of aggregate did not show any Hsp78 signal in the pellet fractions (P.C) (Fig. 2E), indicating that there was no unspecific aggregation of soluble chaperone species that could lead to false positive results. Taken together, the results indicate a stronger substrate affinity for the mutant Hsp78_TR_ compared to the WT protein, consistent with the expected effect of the mutagenesis.

### Co-purification studies reveal a variety of interacting mitochondrial proteins

In order to perform an unbiased identification of potential substrate proteins of Hsp78 in a natural organellar environment we used the tagged WT Hsp78 and mutant Hsp78_TR_ as overexpressed bait proteins in a Ni-metal-affinity based purification procedure. Samples of the co-purification experiments confirmed a successful purification of the Hsp78 proteins, while the outer membrane protein Tom40, serving as internal control for an abundant non-interacting protein, showed no co-purification (Suppl. Fig. 3B), indicating the specificity of our approach. The three different yeast strains expressing un-tagged Hsp78, wild-type tagged Hsp78 and the tagged Hsp78TR mutant were cultured in minimal medium containing light, medium and heavy nitrogen isotope amino acids, respectively. After isolation of mitochondria, organelles were lysed under native conditions and the metal affinity purification was performed. The eluates containing the different Hsp78 forms as well as co-purifying polypeptides were combined (Suppl. Fig. 3C) and subjected to SILAC based quantitative mass spectrometry (Suppl. Fig. 2). Proteins identified in the untagged Hsp78 control sample were defined as background. These background abundance values were substracted from the corresponding signal in the samples containing the respective tagged Hsp78 form as bait to allow a positive identification of potential interacting proteins.

In total 465 proteins were at least once identified by MS in the different co-purification experiments (raw qMS abundance values summarized in Table S2) of which 208 proteins exhibited a positive value above background. To correct for the differences in bait (Hsp78) expression levels, the interaction values were normalized to the qMS abundance of the respective Hsp78 variant (Table S3). To assess the specificity of the approach, we sorted all positively identified proteins according to their assigned cellular localization and plotted them based on their relative abundance values (Fig. 3A). Except for the WT Hsp78 samples at 25°C, the cofractioned proteins were predominantly mitochondrial, indicating the validity of the approach. In most samples more than 90% of the co-fractionating protein amount were mitochondrial, with the majority located in the matrix compartment, as expected. The comparably large amount of non-mitochondrial proteins co-fractioning with Hsp78 at 25°C (about 45% of the total) likely indicated a high probability of non-specific interactions that occur after lysis of mitochondria. In this sample, 41% of the total protein amount was represented solely by the proteins Lsp1 and Pil1, both very abundant proteins and components of the eisosome complex that is associated with the cell membrane. Hence, we suggest that these were purification artefacts. Another outlier protein was the mitochondrial outer membrane protein Om45 that was found in all samples in very high amounts, representing at least 25% up to more than 80% of the cofractionating protein amounts. As Om45 is an integral membrane protein of the outer membrane with its bulk part exposed to the cytosol, it is very unlikely that it represents a potential Hsp78 substrate and was therefore excluded in the subsequent analysis of the quantitative data. In case of the WT form, the proportion of non-mitochondrial proteins decreased strongly during and after the 42°C heat shock treatment, indicating that under stress conditions the interaction specificity switched to the real endogenous substrate polypeptides. Interestingly, the overall binding behaviour of the Hsp78_TR_ mutant at 25°C closely reflected that of the WT form under heat stress. This behaviour likely correlates with the expected increased substrate binding affinity of Hsp78_TR_ already in absence of stress conditions. We also detected some inner membrane proteins binding to both Hsp78 forms with total ratio between 4% and 8%.This suggests that also some inner membrane proteins, likely that are exposed to the matrix compartment, can serve as potential Hsp78 substrates. In addition, the specific effects of the different heat treatments already indicate a broad role of Hsp78 concerning mitochondrial thermal tolerance.

**Fig. 3:**
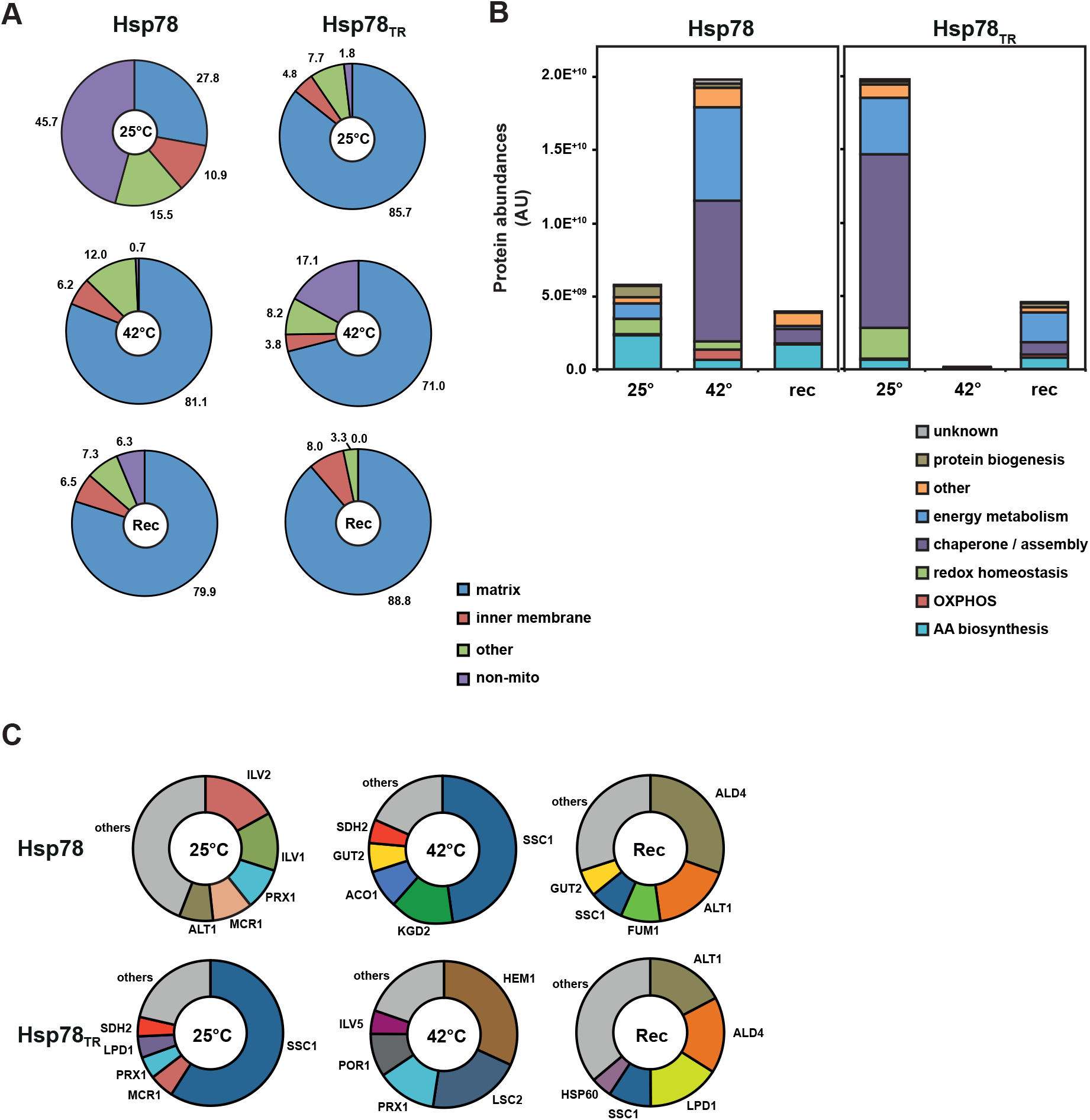
Hsp78 interaction proteome analysis by affinity-tag co-purification. A, Subcellular localization distribution of proteins co-purifying with Hsp78 and Hsp78TR after the indicated three temperature treatments. Protein hits were categorised by UNIPROT annotations. Protein qMS abundances were calculated as normalized average abundance over background from three independent experiments (see Methods section; data taken from Table S3). Relative values for abundance sums for the indicated cellular locations are shown (total sum of all protein abundances was set to 100%). B, Abundance sums for WT and mutant co-purification results sorted by the indicated functional categories. Only mitochondrial proteins are shown. OM45 has been omitted as a likely artefact. C, Relative ratio of the 5 most abundant proteins found under the different conditions with both Hsp78 variants.

In order to assess the relevance of Hsp78 protein interaction for the function of mitochondrial proteins, we assigned the identified Hsp78-interacting mitochondrial proteins to their respective functional categories and sorted them by their signal abundances, reflecting the absolute amounts of Hsp78-bound polypeptides (Fig. 3B). Concerning the total abundance sum of all interacting proteins, we observed a strong increase of protein binding for WT Hsp78 at 42 °C, supporting the hypothesis that Hsp78 is mainly functional under heat stress conditions. During the recovery period, the amount of bound proteins decreased significantly, suggesting that many substrate proteins are released from the chaperone after the heat stress has subsided. However, the pattern of functional categories bound during the recovery was different from that under normal conditions, indicating that certain substrate proteins undergo a prolonged interaction with the chaperone. The most abundant Hsp78-interacting proteins at 42°C mainly fit into the categories ‘chaperones/protein assembly’ and ‘energy metabolism’ (mainly TCA cycle components), while the proteins from the categories ‘redox homeostasis’, ‘OXPOS’ and ‘amino acid (AA) biosynthesis’ were less strongly represented. In comparison, the mutant Hsp78_TR_ showed the highest amount of interacting polypeptides already at 25°C and almost no binding affinity under stress conditions, indicating the effect of the putative substrate trap mutation and correlating with a functional defect at elevated temperatures. Interestingly, the pattern of functional categories between Hsp78_TR_ at 25°C and WT at 42°C was quite similar, suggesting that the mutations did not substantially alter the substrate range of the chaperone. The high amounts of Hsp78-interacting proteins belonging to the clusters ‘energy metabolism’ and ‘chaperones’ likely represent functional clusters of chaperone substrate proteins that are important for the maintenance of key metabolic functions of mitochondria.

Up to 70% of the total protein abundance in the different co-purification samples was composed by a few individual polypeptides, most likely representing the main substrate polypeptides of Hsp78. For WT-Hsp78 at 25°C these included the proteins Ilv2 (17%), Ilv1 (13%) and Alt1 (8%), all enzymes involved in amino acid biosynthesis, as well as Prx1 (9%), Mcr1 (9%), both redox homeostasis, while 50% of the signal was generated by a mixture of less abundant proteins. Switching to 42°C, the matrix chaperone Ssc1 became predominant with almost 50% of the total signal intensity. Moreover Kgd2 (14%), Aco1 (9%), Gut2 (6%) and Sdh2 (5%), all metabolic key enzymes, were very abundant and the remaining 18% consisted of a mixture of proteins. Under recovery conditions, the amount of Ssc1 decreased again to 8% while the abundant metabolic enzyme Ald4 (30%; aldehyde dehydrogenase) became more abundant together with Alt1 (17%), Fum1 (9%; TCA cycle), and Gut2 (6%). The intensity of other proteins increased to 29%. For Hsp78_TR_ the Hsp70-chaperone Ssc1 represented the most abundant interactor with 59% already at 25°C, while Mcr1, Prx1, Lpd1 (a major component of the pyruvate dehydrogenase complex) and Sdh2 were found with around 5% each. At elevated temperatures, the aminolevulinate synthase Hem1 (32%), Lsc2 (21%, TCA cycle), Prx1 (9%), Por1 (9%, an abundant outer membrane protein) and Ilv5 (5%, AA biosynthesis) became more abundant while Ssc1 was strongly decreased. During recovery, Alt1 (17%) and Ald4 (17%) were the most abundant interaction partners (similar to WT), followed by Lpd1 (16%), and the chaperones Ssc1 (9%) and Hsp60 (5%) were the most abundant hits (Fig. 3C).

In order to visualize the behaviour of the protein population and its Hsp78 binding tendency as a whole, log_2_ values of the quotients between sample versus background protein abundances were plotted for the different conditions. For individual proteins a ratio of 1 (log_2_ of 0) and above implies an enrichment of the proteins by the co-purification and hence binding to Hsp78. By plotting this co-fractionation enrichment ratios at 25°C versus those at 42°C we additionally visualized the role of Hsp78 interaction during the transition from normal to stressed conditions. As expected for a chaperone, with WT Hsp78 the majority of interacting polypeptides increased during heat stress. As in this visualization the absolute amount of bound proteins was not relevant, this approach also identified low-abundance proteins that exhibited a strong interaction with Hsp78. For example, the uncharacterized protein YIL045w, a probable minor catalytic subunit of succinate dehydrogenase (SDH), stood out as a major interacting polypeptide at both conditions. Most other proteins were higher enriched at 42°C, notable examples were Aco1 (aconitase), which had been identified already repeatedly in previous publications as a heat-labile mitochondrial enzyme, and Ssq1, another Hsp70 chaperone family member. In addition, the graphical analysis demonstrated again the difference in substrate binding behaviour between the WT protein and the mutant version during the shift from at 25° to 42°C is clearly represented. At 25°C, the mutant bound stronger to these potential substrates than the WT, as the cloud of protein dots accumulate near the horizontal axis while the WT-Hsp78 showed increased protein binding under heat stress conditions, as shown by the accumulation of substrate proteins on the vertical axis (Fig. 4A). Plotting the interaction data under heat stress versus recovery period, we observed a more similar behaviour of both Hsp78 variants (Fig. 4B). Nevertheless, the mutant Hsp78_TR_ revealed a strong binding of some proteins under recovery conditions that were not prominent as substrate polypeptides under heat stress conditions. Examples for this group are the proteins Hem1 (5-aminolevulinate synthase), Mcr1 (NADH-cytochrome b5 reductase), and Lsc2 (subunit of the succinate-CoA ligase). It is conceivable that – depending on the individual structural properties of the respective proteins – an interaction with Hsp78 still persists even after a return to normal conditions in order to support potential refolding reactions.

**Fig. 4:**
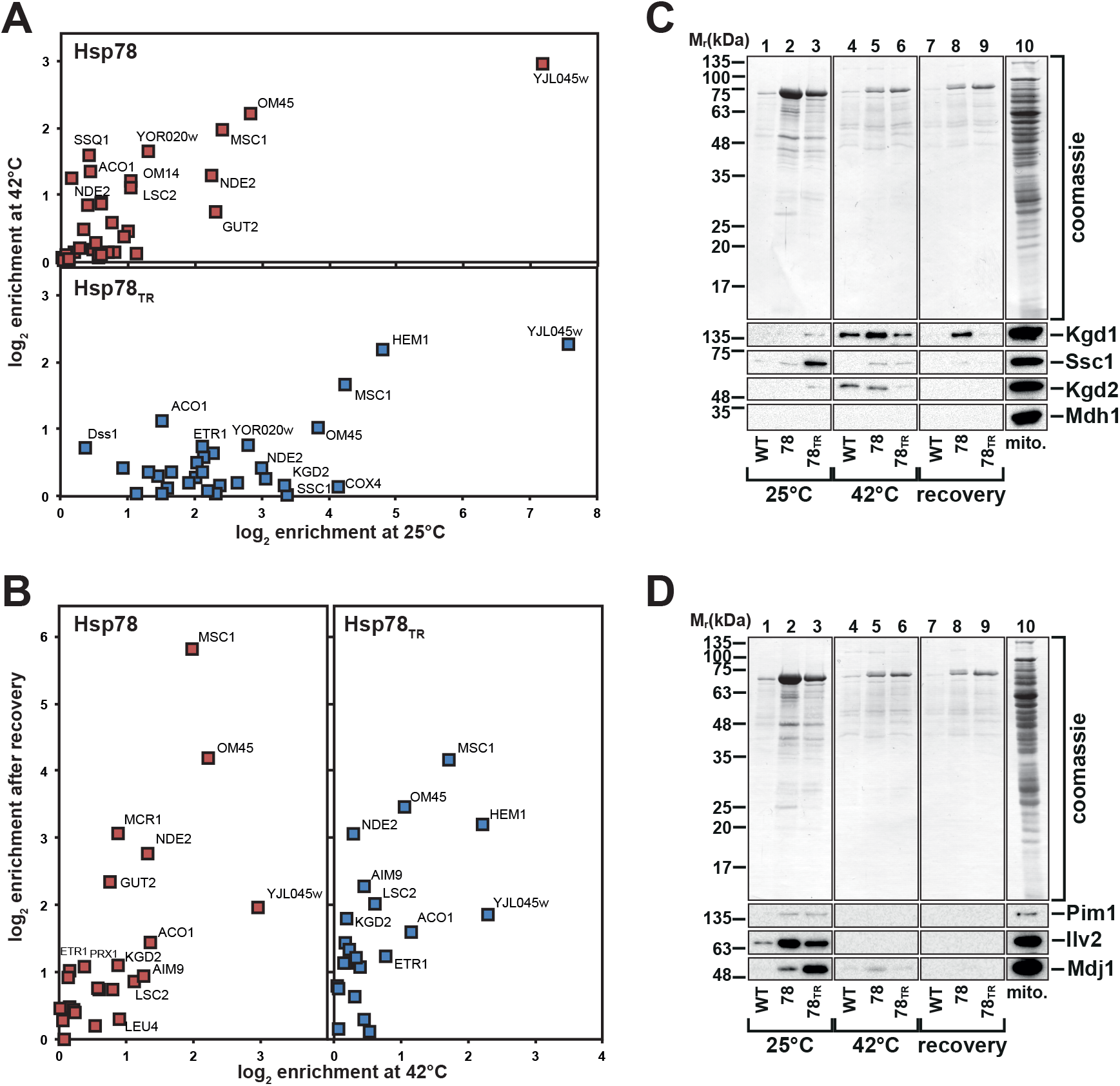
Co-purification experiments reveal different behaviour of Hsp78 and mutant Hsp78TR under various conditions. A, Comparison of substrate binding at normal conditions (25°C) and under heat stress (42°C). Scatter plot for wild-type (blue) and mutant (red) Hsp78 co-purification ratios over background signals (log2 values), expressing enrichment by binding affinity. Ratios are calculated from normalized average qMS abundances of three independent experiments (see Methods section; data taken from Table S3). Proteins that were detected but not enriched over background values were excluded. Strongly enriched proteins are identified. B, Comparison of substrate binding at heat stress normal conditions (42°C) and after 1 h recovery back at 25°C. Data shown as described in A. C and D, Co-purification experiment as described using WT (Hsp78) and mutant variant (Hsp78TR) mitochondria. Eluates were analysed by SDS-PAGE and Western blot using the indicated antisera. As loading control the Coomassie Blue stained membrane is shown. Lane 10 (mito.) shows signals obtained with a corresponding amount of total mitochondria.

We also tested the aggregation behaviour of selected individual proteins by Western blot to confirm our qMS data (Figs. 4C and D). Overall, the results confirm the qMS characterization, exhibiting a very variable interaction pattern at the different temperatures and also between the two forms of Hsp78 used. Both subunits of the ketoglutarate dehydrogenase complex (Kdh1 and Kdh2), exhibited a strong binding to Hsp78, in particular to the WT form, confirming their sensitivity of heat-induced aggregation (Fig. 4C). The amino acid biosynthesis enzyme Ilv2, showed its strongest interaction already under normal temperatures, indicating a potential conformational instability that requires the chaperone activity of Hsp78 already under normal environmental conditions (Fig. 4D). The members of the mitochondrial Hsp70 system, Ssc1 and Mdj1 (the corresponding member of the DnaJ co-chaperone family) showed a high interaction affinity in particular with the Hsp78_TR_ mutant. The metabolic enzyme Mdh1 was served as a control for a stable soluble protein and did not co-purify (Fig. 4C).

### Identification of aggregation-prone and actively disaggregated proteins

To obtain a global overview about mitochondrial protein aggregation and disaggregation reactions, we performed an additional quantitative proteomic characterization of the general *in organello* aggregation processes, using SILAC-assisted mass spectrometry. We determined protein abundances in the high-speed centrifugation pellet fractions of isolated detergent-lysed mitochondria to recover insoluble proteins as putative aggregates (prepared as described above) under different conditions. As control, mitochondria were kept at 25 °C while aggregation was induced by 30 min heat treatment at 42 °C. A potential disaggregation of polypeptides from aggregates was also assayed after a recovery incubation at 25 °C for 1 h after the heat treatment. To assess the impact of Hsp78 activity on these processes we again used the Hsp78 WT and substrate-trap variants described above. We detected in total 1199 different proteins in the pellets fractions (Table S4), of which 470 were localized to mitochondria as assigned by the UniProt database, comprising on average about 96% of the total pellet protein abundance. From the 379 mitochondria-associated proteins identified at least once under every condition and in each Hsp78 variant (Table S5), we categorized 93 (about 25%) as strongly aggregating proteins, meaning they showed a more than 4 times higher relative abundance in the pellet after 42 °C compared to the 25 °C pellet in mitochondria expressing WT Hsp78 (Fig. 5A and Table S5). Under the 5 most abundant proteins in the aggregate pellet were Aco1, Lsc2 (both citrate cycle), Hsp60 (chaperone), Ilv5, Ilv2 (both aa biosynthesis) and Gut2 (energy metabolism). In contrast to the matrix chaperonin Hsp60, Hsp78 itself was not identified in any of the aggregate pellets, indicating that it retains its solubility even during heat stress conditions, also indicating that it does not stably interact with aggregating polyeptides. The main chaperone of the matrix compartment, Ssc1 or mtHsp70, was highly abundant in the heat stress pellet fraction but also relatively strongly represented already under normal conditions and therefore barely did not meet the criteria for a strong aggregator. Of this list of strong aggregators, many were already identified above as potential Hsp78-interacting proteins, in particular the already mentioned Aco1, Lsc2 and Gut2. In addition, the identified interactors Mcr1, Prx1. Fum1, Alt1, Sdh2 and Hem1 exhibited also a high aggregation ratio. When analyzing the pellet composition after the recovery period, we observed in general no significant resolubilization of high-abundance aggregating polypeptides, suggesting that the effects of a high protein amount in the aggregate pellet together with a strong aggregation propensity cannot be reversed by a disaggregation system during the reaction time of 240 min. However, some low-abundance proteins with significant aggregation propensities showed significant disaggregation (Fig. 5B, more than 50% reduction in the pellet), for example Afg1 (unknown function), Nfu1 (iron sulphur cluster biosynthesis), Mef2 (translation), Leu9 (aa biosynthesis) and Coq9 (ubiquinone biosynthesis). These actually disaggregating polypeptides were on average at least 10x less abundant in the aggregate pellet than those that were not solubilized. Also a few other low-abundance proteins showed a high disaggregation potential: Trr2 (redox reactions), Cox2, Atp11, Cir2 (all involved in respiration), Mrs2, Ala1, Vas1 (protein expression/translation), as well as Tim23 (protein transport) and Tum1 (sulfur metabolism). Considering a potential general negative effect of the Hsp78_TR_ mutation on the aggregation reactions, we observed that the group of high-abundance aggregating proteins were barely affected, meaning the composition of the aggregate pellets of WT Hsp78 and Hsp78_TR_ was remarkably similar, both directly after the heat stress as well as after the recovery period (Fig. 6). Only a few proteins, for example Adh4 (mitochondrial alcohol dehydrogenase), Rip1 (iron-sulphur protein of respiratory complex III) and Adk2 (GTP:AMPphosphotransferase) show significantly higher aggregation in Hsp78_TR_ than in WT at 42°C as well as after the recovery. It is therefore likely that these proteins are particularly dependent on Hsp78 activity for maintaining their solubility. However, this picture was different for the composition of the aggregate pellet at 25 °C. Here many proteins that were not strongly represented in the pellet of WT mitochondria, in particular with generally lower abundances, showed a higher aggregation rate in the Hsp78_TR_ mutant (Fig. 6). This indicated that a distinct group of proteins is dependent on at least some assistance by Hsp78 to maintain their full solubility even under normal growth conditions.

**Fig. 5:**
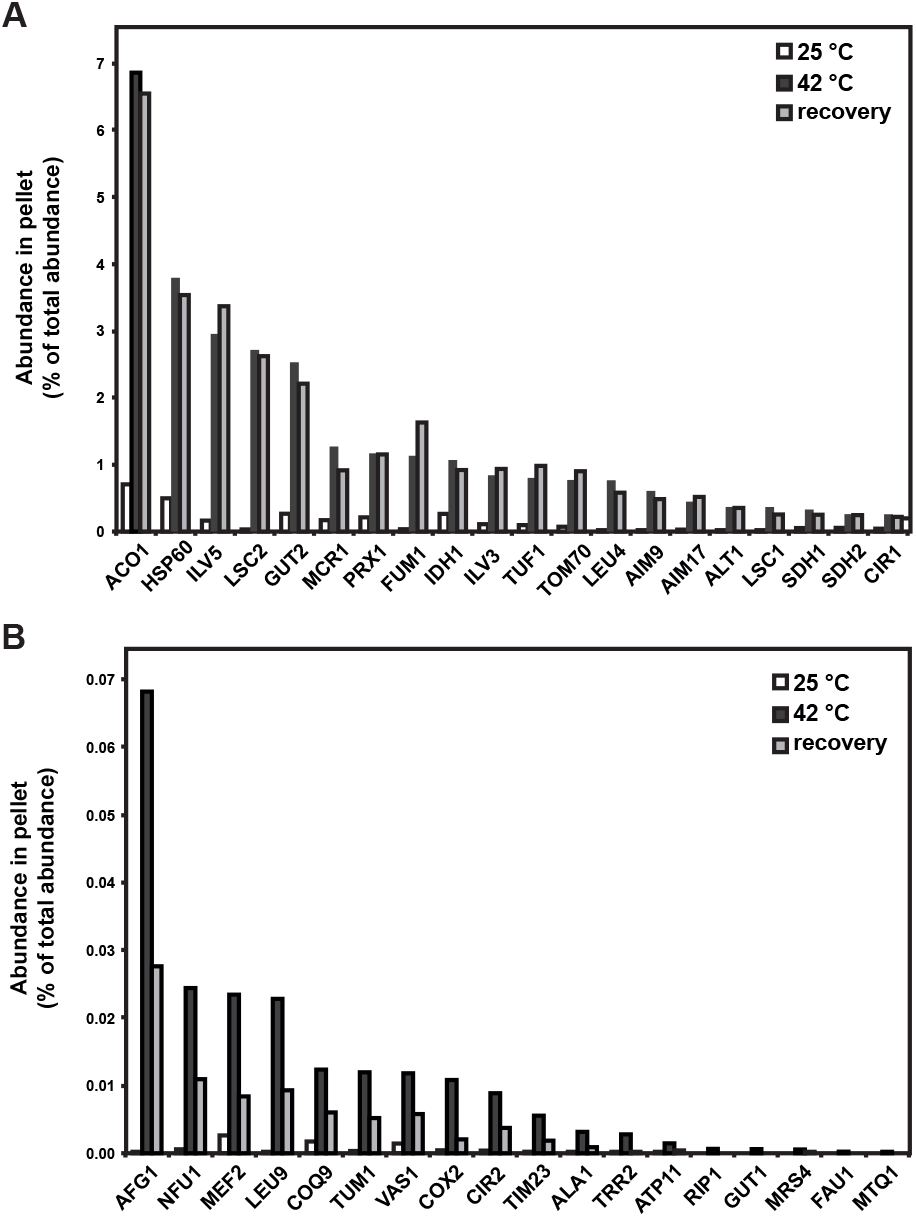
Identification of aggregation-prone mitochondrial proteins. A, Relative abundance ratio in the aggregate pellets of Hsp78 and Hsp78TR-expressing mitochondria incubated under the indicated temperature treatments (total sum of all protein abundances in the respective pellets was set to 100%). Shown are the 20 most abundant proteins in the pellets after 42°C. Protein abundances were calculated as normalized average abundance over background from three independent experiments (see Methods section; data taken from Table S4). B, Same dataset as in A. Shown are the 20 proteins with the highest disaggregation ratio (difference between abundance after 42°C heat stress and after 1 h recovery).

**Fig. 6:**
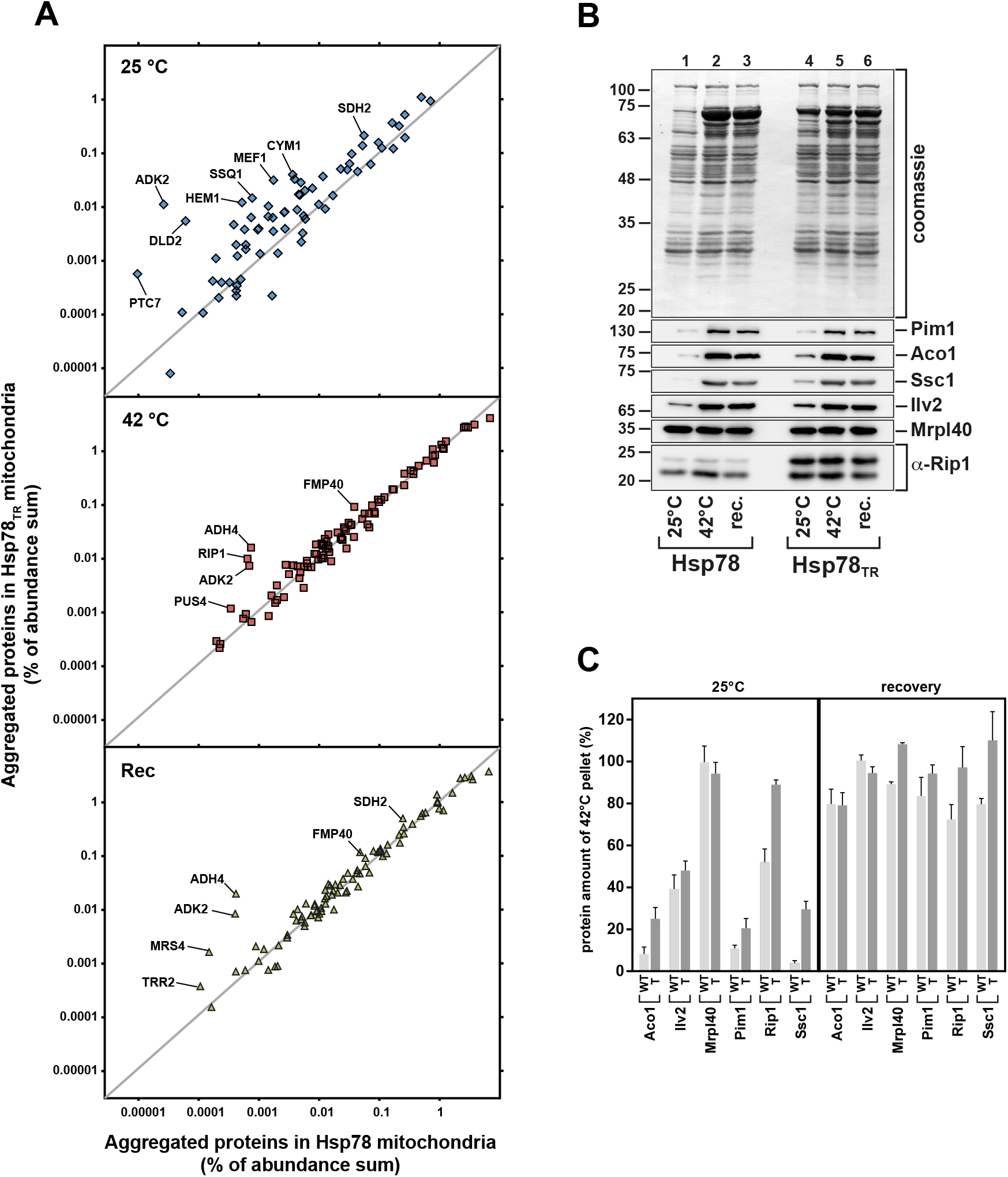
Proteomic aggregation and recovery assay identifies aggregation-prone proteins and reveals aggregated proteins that are re-solubilised by Hsp78 function. A, Scatter plots comparing relative abundances of proteins in aggregated pellets for WT (Hsp78) and mutant (Hsp78TR) expressing mitochondria under the indicated temperature-treatments. Dots near the diagonal represent proteins that do not differ between the two Hsp78 variants. Protein candidates with higher aggregation tendency in Hsp78TR are identified (data taken from Table S4). B, Western blot analysis of aggregated pellets and control Coomassie Blue staining derived from mitochondria expressing Hsp78 or Hsp78TR treated with the indicated temperatures. Membranes were decorated with the indicated antisera. C, Quantification of Western blot characterization of aggregate pellets mitochondria expressing Hsp78 (WT) or Hsp78TR (TR) as shown in B (n=3). Values are calculated for 25°C and recovery conditions relative to the signal intensity at 42°C (100%).

For some proteins with a high aggregation tendency, including Pim1, Aco1, Ssc1, Ilv2 and Rip1, we also confirmed their aggregation behaviour by a Western blot analysis. All became insoluble after 20 min of 42°C treatment and were therefore enriched in the pellet fraction (Fig. 6B and C). Potential substrates were present with higher signal intensities in the pellet fraction of the Hsp78_TR_ mutant at 25°C, which provides further evidence to a trapping effect in the mutant. Interestingly, Rip1 showed both an increased aggregation and a decreased resolubilisation rate in the mutant mitochondria, indicating an Hsp78 involvement and directly confirming the qMS results (Fig. 5B). The mitochondrial ribosomal complex protein Mrpl40 was expected to pellet under the chosen experimental settings and served as a positive sedimentation control (Fig. 6B).

With a combined number of 271 proteins, around 25% of all mitochondrial proteins were found to interact in some fashion with Hsp78. 41 proteins were observed in both mass spectrometry approaches representing key proteins from different cell-functional groups. With the help of the internet-based proteome analysis tool STRING (functional protein association networks; string-db.org), we generated a interaction network for all putative Hsp78 substrate proteins, including basal providing also functional information for them (Fig. 7). Proteins that do not have listed interaction partners inside the network are not shown. The network illustrates an accumulation of proteins in functional nodes representing mitochondrial translation, TCA cycle, oxidative phosphorylation, ATP synthase complex assembly, transport, amino acid biosynthesis, iron-sulphur complex binding and chaperone-mediated protein complex assembly. Furthermore, we can see a high degree of connectivity within the protein groups for TCA cycle and especially for mitochondrial translation. The cluster for translation includes many components of the small and large ribosomal subunits, which are known to interact closely. Taken together, Hsp78 seemed to represent a key regulator of protein solubility after heat exposition that is not specialised on specific functional pathways but exhibited the behaviour of a general chaperone with no evident substrate recognition properties.

**Fig. 7:**
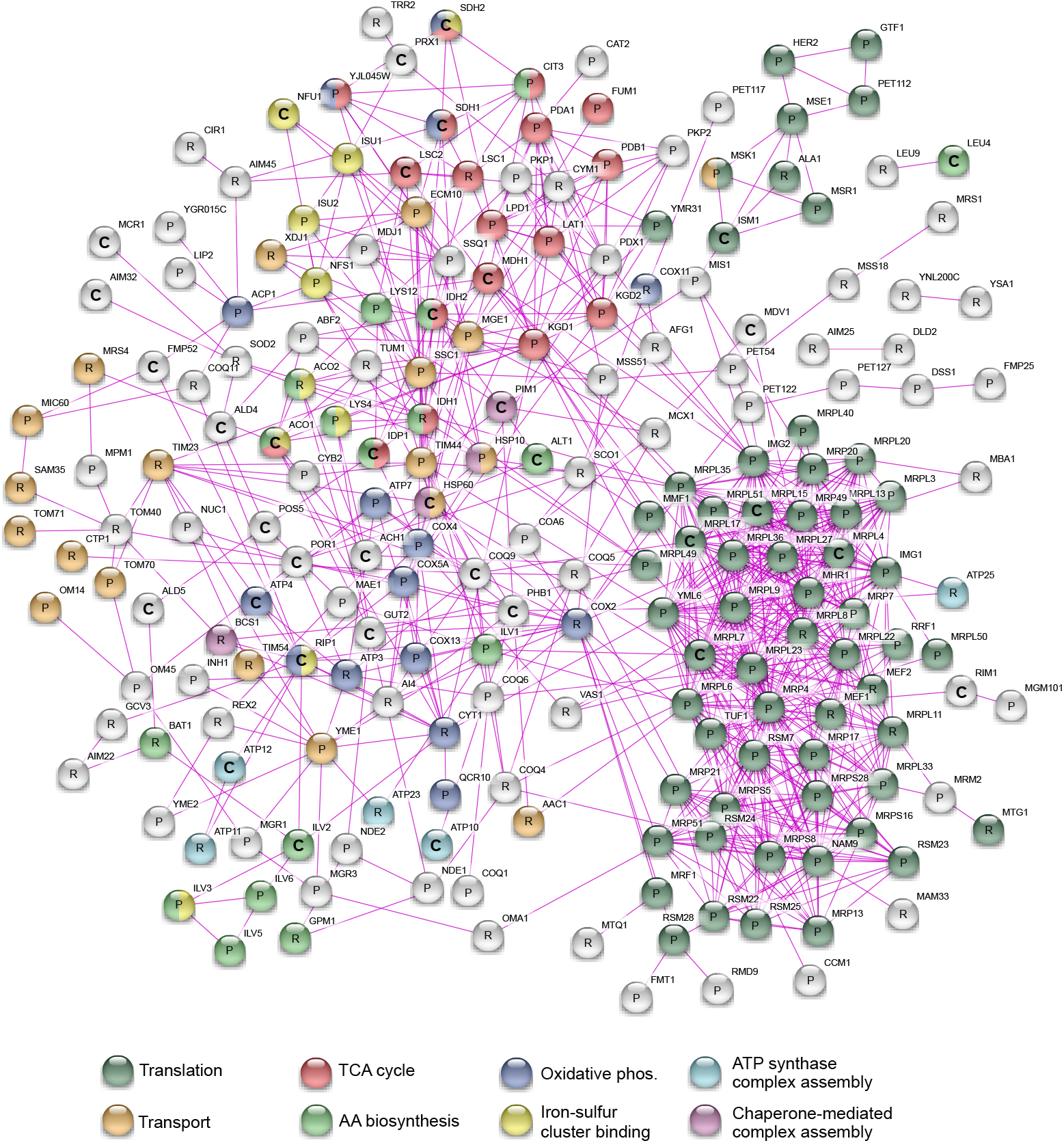
Important mitochondrial functional clusters are affected by Hsp78 interaction. A, Interaction network of Hsp78-affected proteins based on STRING (functional protein association network) data analysis, using qMS results from Hsp78 co-purification mitochondrial aggregation and recovery experiments. Illustrated are proteins as nodes found in either co-purification (P), in aggregation and recovery pellets (R) or in both approaches (C). The involvement of proteins in mitochondrial functions (GO annotation) is shown by node colour. Protein-protein interactions are displayed for experimentally proofed cases according to database. With 271 nodes the statistically expected number of edges should be 213 while the dataset shows 863 edges with an average node degree of 6,37. The PPI enrichment p-value is lower than 1.0^−16^.

## Discussion

With the goal to establish a comprehensive picture of the mitochondrial protein aggregation behaviour under heat stress we performed two independent quantitative proteome analysis approaches. We focused on the function of the mitochondrial Hsp100/ClpB family member Hsp78, as ClpB-related proteins have been established as the major chaperone system responsible for prevention of temperature-induced protein damage (3). Although we previously identified a small number of aggregating mitochondrial proteins (31), we reasoned that only a proteomic characterization of protein aggregation will provide an understanding of a potential substrate hierarchy for Hsp78 and provide fundamental knowledge about mitochondrial protein behaviour under stress. The first approach utilized an affinity-based co-purification with Hsp78 as bait to identify interacting proteins that represent potential chaperone substrates. The second approach was based on an overall analysis of intramitochondrial aggregation and disaggregation processes in presence or absence of active Hsp78. Aggregation reactions were performed based on an already established assay using isolated intact organelles to focus on mitochondrial proteins (27,31). As many chaperones have only a limited affinity to their substrate proteins, we tried to increase the chance of substrate identification by constructing a substrate-trap form of Hsp78 that carried point mutations in the Walker B motifs of the ATPase domains. Similar setups have already been used for Hsp100 substrate identification in bacteria (24,32). Previous studies on Hsp78 as well as its bacterial homolog ClpB either just used *in vitro* approaches using selected model substrates or focussed solely on an identification of interacting polypeptides via MS. In comparison, we aimed on a combination of two strategies, the identification of potential substrates and a concomitant characterization of the overall aggregation propensity of the mitochondrial proteome, in both cases utilizing a quantitative proteome analysis approach in the WT and substrate trap background.

The biochemical characterization of the Hsp78TR mutant demonstrated its suitability for the chosen approach. Similar to the corresponding ClpB mutant in *E. coli* (33), the Hsp78_TR_ mutant a strongly reduced ATPase activity. The new Hsp78_TR_ mutant protein essentially reproduced the effects of a *hsp78Δ* deletion mutant, showing strongly reduced disaggregation potential for imported reporter constructs (27) and a decline of acquired thermotolerance under conditions that require the metabolic activity of mitochondria (34). The formation of hexameric chaperone complexes is a typical hallmark of Hsp100 family proteins that is closely related to their functional activity (35). In line with previous published data, both Hsp78 variants assembled into higher oligomeric states favoring dimers or trimers, which in turn associate to hexamers (17,18). We found that Hsp78_TR_ exhibited a slightly higher complex stability and also showed complexes with higher molecular weight than expected for hexamers. This observation might indicate that substrate release is reduced or delayed as the full dynamics of complex dynamics is required for the substrate interaction and disaggregation activity of ClpB-type chaperones (36). This would also be consistent with the reduced ability of Hsp78_TR_ to hydrolyze ATP, which is also correlated to the dynamics of the enzyme complex (17,18). Hsp78_TR_ also exhibited a slightly higher binding affinity to aggregated polypeptides *in vitro* compared to the WT protein, indicating its suitability for substrate identification studies.

The tagged Hsp78 constructs were well suited for affinity co-purification experiments and the mutant forms exhibited a direct influence on substrate/aggregate binding as well as effects on the overall aggregation propensity of the mitochondrial proteome. Our approach yielded a high number of proteins identified in the qMS analysis that served as new interaction and/or aggregation candidates. Data matching with the available information in the protein database revealed that the large majority of hits represented mitochondrial proteins, in most cases localized to the matrix compartment. In addition to the matrix components also proteins from the inner membrane with segments exposed to the inside can be expected as potential Hsp78 substrates, demonstrating the validity of the used experimental approach. However, a few proteins from other cellular compartments were also identified. Some of them may result from cross contaminations in particular from ER proteins or other vesicular components that are well known to occur in fractions representing isolated mitochondria. The identification of such proteins in the highly sensitive qMS experiments may reflect minor co-purification artefacts after lysis of the mitochondria. In our further analysis such potential artefacts were identified by a comparison to the control sample containing a Hsp78 form lacking the tag and were discounted in the data analysis. The relative abundance of substrate proteins from the mitochondrial matrix was the highest for Hsp78_TR_ at standard conditions, confirming a direct and more stable substrate interaction resulting from the ATP-hydrolysis alteration inside the Walker B motifs.

Quantitative mass spectroscopy as was performed in our analysis allows not only the identification of potential interactors but also to determine the relative amounts of proteins bound. This provides important information about the functional relevance of the different protein hits and an estimation of the absolute substrate-flow through the chaperone system under different conditions. The total abundance sum of bound polypeptides showed a very strong increase in WT Hsp78 under heat-shock conditions, indicating its role as a heat-stress protective chaperone. The identities of interacting proteins also strongly changed during heat shock, revealing specific substrates or partner proteins. Concerning the overall interaction with proteins, WT and mutant Hsp78_TR_ behaved quite similar. However, Hsp78_TR_ showed a high binding affinity already under normal conditions, correlating most likely with the substrate-trap mutations of the ATPase domain. We identified proteins from many different functional categories. In particular, enzymes of the energy metabolism (TCA cycle and ATP synthesis) as well as chaperone proteins were prominent Hsp78-interacting proteins, supporting the observation that Hsp78 performs an important role in the maintenance of mitochondrial functions during heat stress. As expected, the overall amount of bound polypeptides was reduced in the recovery period but many proteins were not completely released, probable representing substrates that were still in the process of refolding or disaggregation. Interestingly, proteins from certain functional groups differed in the binding behaviour to Hsp78 during heat stress and recovery, respectively. The strong binding to components of the protein biosynthesis machinery and to chaperones during heat stress was almost absent again in the recovery period while the binding to components of the energy metabolism was retained longer even after reduction of the temperature stress. This observation may coincide with the notion that maintenance of the protein biosynthesis machinery, consisting of ribosomal proteins and their respective cofactors as well as chaperone proteins, is the primary task of the cellular stress protection system. In this line, a proteomic characterization of heat stress-related aggregation in mammalian mitochondria demonstrated a fast shut-off of protein translation as a dominant and primary reaction via the inactivation of the translation factor TUFM (11). It is known from other studies that the mitochondrial translation together with DNA replication is strongly impaired after heat stress (23,37,38). Hence Hsp78 activity is of high relevance for recovery of the intra-mitochondrial protein synthesis (34). Especially the identification of a high number of ribosomal proteins and other components of the mitochondrial translational machinery as substrates is reminiscent of a phenomenon described in the cytosol. As a typical reaction upon heat stress, the formation of cytoplasmic stress granules (SGs) or processing bodies was observed. These granules represent dynamic structures that can promote a rapid reactivation of protein biosynthesis after heat –stress has subsided (39). For example HSF-1, the key transcriptional factor for the heat-induced cellular response to temperature stress also forms SGs (40). Hence, a fast regulation of translational processes in response to stress conditions seems to be a primary task for all types of cell-protective mechanisms. The individual polypeptides that represented the most abundant binding proteins are most likely conformationally semi-stabile proteins that tend to loose their native folding state under elevated temperatures. However, not much information is available about the conformational stability of mitochondrial proteins per se, with the potential exception of Adk2, which has been characterized as a highly temperature-unstable matrix protein (41). Indeed, ADK2 belonged to the few proteins that exhibited a low disaggregation efficiency. At least some of the identified proteins, like Kgd2, Aco1 or Ilv2, were already identified as aggregation-prone polypeptides before (31) and therefore represent major Hsp78 substrates. The main mitochondrial Hsp70, Ssc1, is playing a special role in this case, as the highly abundant chaperone was already found to interact with Hsp78 (27). An interaction of Hsp78 with Ssc1 seems to be critical since it was shown that Hsp70s are able to stimulate the ATPase activity of Hsp100 family proteins (42). Therefore it is likely that under heat stress conditions, Ssc1 is also an interaction partner of Hsp78 and not only a substrate itself. Due to the ATPase mutation, this functional interaction with Ssc1 was already apparent for Hsp78_TR_ under normal conditions.

In addition to the Hsp78 interaction data we added a second proteomic data set, aiming on the quantitative characterization of Hsp78 influence on actual aggregation and disaggregation processes inside mitochondria. Again, we used the impaired functionality of Hsp78_TR_ as a tool to monitor *in organello* disaggregation in comparison to WT. Comparing individual protein abundances in the aggregate pellet fraction, we were able to define which proteins aggregate and which are disaggregated by Hsp78 and may fail to recover in the Hsp78_TR_ mutant. In a recent proteomic characterization of yeast protein thermal stability it was proposed that bigger and more flexible proteins that provide more protein-protein interaction sites have a higher tendency for aggregation (43). Our dataset did not confirm the presumption for protein size, but revealed a high number of aggregation-prone proteins that exhibit specific co-factor interactions. These candidate proteins include ion binding (95 hits), nucleotide binding (60 hits), coenzyme binding (18 hits) or iron-sulfur-cluster binding (13 hits). As they exhibit the necessary flexibility that allows a structural adaptation for the co-factor binding, the likelihood of a thermal instability leading to increased aggregation is certainly feasible. We were able to directly confirm the aggregation of some selected proteins and also the main mitochondrial Hsp70-type chaperone Ssc1, but not for its cochaperones Mge1 or Mdj1, which were previously found in aggregates (12). This might be due to the higher temperature of 48°C and the longer heat exposition used in the former study. Moreover we identified many proteins that were also found in a study on bacterial aggregation (bacterial homologs in brackets), such as Put2 (PutA), Kgd1 (SucA), Pda1 (AceE), Adh3 (AdhE), Aco1 (AcnB), Pim1 (Lon), Mae1 (SfcA), Sdh1 (SdhA), Lat1 (AceF) or Fum1 (FumA) among others (8), indicating an evolutionary conserved stress sensitivity of bacteria and mitochondria. Although not much information about aggregation-prone proteins is available for mammalian mitochondria, we also could confirm a strong aggregation tendency for the yeast elongation-factor Tuf1, the homolog of the mammalian Tufm, which was found to be one of the strongest aggregating proteins in HeLa cells (11). Many of the strongly aggregating polypeptides were also found as prominent Hsp78-interacting proteins, confirming the general chaperone role of Hsp78 during aggregation processes. However, we also observed that the highly abundant polypeptides in the aggregate pellet did not exhibit significant disaggregation rates during the recovery period. The proteins that were found to be disaggregated were of lower abundance and also not prominently represented as Hsp78-interacting polypeptides. This indicates that the disaggregation capacity of Hsp78, which itself is not an abundant mitochondrial protein, is limited to polypeptides with low expression levels. This does not exclude that also more prominent proteins can also disaggregated, but the impact on their high amounts in the aggregate pellet is small. As it is known that proteins disaggregated by Hsp78 can be directly degraded by the mitochondrial protease Pim1 (44), they might share a common substrate pool. Looking at all strongly abundant proteins identified in our binding studies it turns out that most of them (except Sdh2, Gut2, Hem1 and Fum1) have been previously identified as substrates for Pim1 (45). We were able to confirm Ilv5 and Lys4 by co-purification and Aco1, Ilv2 and Lsc2 by both techniques. This finding is not surprising since both enzymes, Hsp78 and Pim1, belong to the AAA+ protein family and may share a similar mechanism of substrate protein interaction.

Taken together, we could show that a high percentage of mitochondrial proteins depend on Hsp78 action during thermal stress conditions. Hence, Hsp78 represents the major protein disaggregase on a proteome level in mitochondria, rebooting processes required for normal mitochondrial homeostasis, like intra-mitochondrial translation, as well as all other crucial functions, like energy metabolism. The establishment of an *in vivo* aggregation and recovery assay, also under other stress conditions, will be of future interest to address the balance between processes like chaperone activation, protein sequestration and mitophagy, which all contribute to mitochondrial and cellular fitness. It should be noted that in addition to the activity of the diverse chaperones also other aggregation protective processes occur in cells and in mitochondria. For instance, a recent study from our group revealed an intramitochondrial aggregation protective pathway that restricts organellar damage by the sequestration of aggregated polypeptides in inert deposition sites called IMiQ (46). Together with the enzymatic PQC system, this process also could contribute to maintain general organellar integrity by relieving the chaperone system from the burden of unfolded and aggregated polypeptides (47). Future work should also focus on revealing the equivalent components in mammalian systems to assert the disaggregation capacity in higher organisms. Better information about the dynamics of protein aggregation and disaggregation in mitochondria will provide important insights into general pathological patterns of diseases related to mitochondrial dysfunction and will therefore help to find new approaches for the stabilization of cellular protein homeostasis under stress conditions.

## Supporting information

Yeast strains and plasmids

Cofractionation qMS data

Cofractionation analysis

Aggregation qMS data

Aggregation analysis

## Abbreviations

BN: blue-native
HSP: heat shock protein
m: mature form
mt: mitochondrial
p: precursor form
PAGE: polyacrylamide gel electrophoresis
PQC: protein quality control
qMS: quantitative mass spectrometry
SDS: sodium dodecyl sulphate
SEM: standard error of the means
SILAC: stable isotope labelling by amino acids in culture
WT: wild-type.

## Acknowledgements

This work was supported by the Deutsche Forschungsgemeinschaft (Grant No. VO 657/5-2 to W.V. and Projektnummer 174793735).

## Author Contributions

W. J. and W. V. conceptualization; W. J., G. C. and M. S. data curation; W. J. and W. V. formal analysis; W. V. project administration; W. J., M. S. and W. V. resources; W. V. supervision; W. J. and M. S. validation; W. J. and W. V. writing—original draft; W. J., M. S. and W. V. writing—review and editing.

