## Supplementary material for "Hsp78-interaction proteome highlights chaperone role during protein aggregation in mitochondria under heat stress": Yeast strains and plasmids

**Table S1.**

| **Strains** | **Genotype** | **Source** |
|---|---|---|
| Y03617-ES | MATa, *his3Δ1, leu2Δ0, met15Δ0, ura3Δ0, hsp78Δ::kanMX4* | Euroscarf |
| Y13617-ES | MATa, *his3Δ1, leu2Δ0, lys2Δ0, ura3Δ0, hsp78Δ::kanMX4* | Euroscarf |

| **Plasmids** | **Description** | **Reference** |
|---|---|---|
| pHSP78 | AmpR, *URA3*, 2μ, pGAL1/10-*HSP78* in pYES2 | (1) |
| pKR09 | AmpR, *URA3*, 2μ, pGAL1/10-*HSP78*-6xHis in pYES2 | K. Röttgers |
| pKR11 | AmpR, $b_2$(107)Δ-$DHFR_{ds}$ in pUHE24 | (1) |
| pTB10 | AmpR, *LEU2*, CEN, Hsp78 ORF in pRS315 | T. Bender |
| pTB25 | KanR, *HSP78*(M385D)-6x-His, in pET28b | T. Bender |
| pWJ01 | KanR, *HSP78*(E216Q/M385D)-6x-His in pET28b | This work |
| pWJ02 | KanR, *HSP78*(E216Q/E614Q/M385D)-6xHis in pET28b | This work |
| pWJ04 | AmpR, *LEU2*, CEN, *HSP78*(E216Q/E614Q/M385D)-6x-His in pRS315 | This work |
| pWJ05 | AmpR, *LEU2*, CEN, *HSP78*(E216Q/E614Q)-6xHis in pRS315 | This work |
| pWJ07 | AmpR, *URA3*, 2μ, pGAL1/10-*HSP78*(E216Q/E614Q)-6xHis in pYES2 | This work |

1. von Janowsky, B., Major, T., Knapp, K., and Voos, W. (2006) The disaggregation activity of the mitochondrial ClpB homolog Hsp78 maintains Hsp70 function during heat stress. *J. Mol. Biol.* **357**, 793-807
